# “Toki”, a light low-cost video system for seabed research: performance and precision of Tehuelche scallop (*Aequipecten tehuelchus*) survey estimates in San José Gulf, Argentina

**DOI:** 10.1101/2024.04.21.590398

**Authors:** G. Trobbiani, L. N. Getino Mamet, A. Irigoyen, A. M. Parma

## Abstract

Studies of benthic abundance and biodiversity are essential for fisheries management and environmental impact assessments. In the case of commercial species, such as Tehuelche scallop (*Aequipecten tehuelchus*), accurate information on the stocks and their densities is crucial. The standard survey method used to estimate abundance of Tehuelche scallops in San José Gulf, Argentina Patagonia, based on visual counts obtained by divers is labor-intensive and limited to diving depths. This study introduces a light low-cost video remote system, named “Toki”, for monitoring scallop beds from small boats.

Toki consists of a steel pyramidal structure that houses two cameras, capable of capturing georeferenced high-definition images. The performance of Toki for estimating the density and size composition of scallops was evaluated. The results demonstrate that Toki provides efficient and accurate density estimates, with variations that can be attributed to observer experience and environmental factors such as bottom algal cover. The size frequency distributions obtained from Toki aligned well with those from hand collection, which allows such measurement for beds beyond diveable depths.

Toki offers advantages in terms of spatial coverage and permanent data records. While there are some limitations, such as processing time for image analysis, future developments based on artificial intelligence can overcome these challenges. Toki has the potential to improve the monitoring of scallop beds and data collection.

**Highlights:**

- Toki offers a low-cost solution for scallops beds monitoring.
- Toki efficiently estimates scallops density within beds.
- Size-frequency distributions obtained from remote videos were accurate.
- Toki contributes to overcoming depth limitations of dive-based scallop surveys.

## 1. Introduction

Benthic abundance and biodiversity studies encompass diverse aims, such as species inventories, stock assessments, habitat characterization and environmental impact assessments (Eleftheriou, 2013; Flannery and Przeslawski, 2015). In species of commercial relevance, such as scallops, accurate information about the exploited stock (e.g., spatial distribution, biomass, size composition) is key for fisheries management (Orensanz *et al*., 2016). Scallop stocks can be assessed by a range of methods that could involve either direct sampling of grounds and beds or indirect methods based on modelling the depletion induced by fishing (Orensanz *et al*., 2016). Among the direct methods, visual counting of scallops along transects carried out by divers has been used in surveys of small-scale fisheries (Amoroso *et al*., 2011; McGarvey *et al*., 2008; Rosenkranz and Byersdorfer, 2004; Stokebury and Himmelman, 1993). The limitations of diving (effort, maximum diving depth, safety issues) together with recent technological advances have motivated the incorporation of remote observations to collect abundance data, especially using video cameras (e.g., towed cameras, drop cameras, remotely operated vehicles, autonomous underwater vehicles) (Batter *et al*., 2021, Fields *et al*., 2019, Stokesbury and Bethoney, 2020, Singh *et al*., 2014, Flannery and Przeslawski, 2015; Goshima and Fujiwara, 1994). Such technologies have demonstrated efficiency for collecting georeferenced data in a wider depth range than diving, making it possible to cover larger areas with less effort and greater security (Bethoney *et al*., 2019; Conan and Maynard, 1987).

The Tehuelche scallop, *Aequipecten tehuelchus*, is a pectinid species distributed in shallow waters of the Southwestern Atlantic Ocean (23-45° S), with commercial relevance in the northern Patagonia gulfs, Argentina (Orensanz *et al*., 2007; Soria *et al*., 2016). New cohorts of juveniles settle each year during the summer and recruit to the fishery when they reach a legal size of 60 mm of shell height, after a year and a half of life (Orensanz, 1986; Ciocco, 1991, 1992a; b). Growth slows down after the second year of life; scallops reach a maximum height close to 100 mm and live up to 11 years; however, specimens over 90 mm and 8 years old are uncommon (Ciocco *et al*., 2006; Soria *et al*., 2016). Spatially, the individuals concentrate in beds, with densities more than ∼25 scallops/m^2^ (Amoroso, 2012; Ciocco *et al*., 2006; Fiorda and Parma, 2015; Parma *et al*., 2008; Soria *et al*., 2017). The scallop beds are mostly composed of single cohorts, but a few age classes can also be found in the same bed. Usually, the 1+ age class can be differentiated by size (30-60 mm), but modes corresponding to older scallops overlap.

The scallop beds support small-scale shellfish fisheries in San Matías (SMG) and San José (SJG) gulfs (Elías *et al*., 2009; Sánchez-Carnero *et al*., 2022). In SJG, this activity has been carried out for close to 50 years by hand-collection using hookah-diving equipment as artisanal or industrial dredges were banned in 1976 (Orensanz *et al*., 2007). Hookah diving and hand-collection is selective, being conducted from small boats (maximum of 9.9 m in length) that generally have two divers (Elías *et al*., 2009). Annual scallop landings currently fluctuate around 200 t, down from a maximum of close to 1200 t reached in 2006 (Getino Mamet *et al*., 2021). The resource has been regulated in a participatory way by the provincial fisheries administration in consultation with artisanal fishermen and scientists (Orensanz *et al*., 2007). Main regulatory measures include a limited entry (maximum of 21 permit holders), a size limit of 60 mm of shell height, a reproductive closure (December-March) and a Total Allowable Catch (TAC) equally divided among permit holders (Orensanz *et al*., 2007; Cinti *et al*., 2011; Orensanz *et al*., 2013). The TAC is calculated based on a survey of the stock (Orensanz *et al*., 2013), performed using a diving-based visual census to estimate the location, biomass, and size structure of scallop beds (Soria *et al*., 2017). As is typical of scallops, annual recruitments are highly variable which requires that the TACs be adjusted annually, considering the biomass that is above and below the legal commercial size (60 mm).

While the scallop survey has been able to track trends in abundance and location of scallop grounds over time, it has three main limitations. First, visual counts in dense areas (i.e., > few individual/m^2^) are imprecise (Amoroso *et al*., 2010, 2011). Second, the method demands a huge effort in terms of diving hours, which requires coordination among various teams to achieve an acceptable coverage and sample size. Third, significant scallop beds have been detected at depths beyond the diving range (> 25 m) that cannot be assessed by the regular survey method (Fiorda *et al*., 2012). The scallop survey was performed quasi-annually from 2001 to 2017, when surveys were discontinued (Soria *et al*., 2017) due to lack of funding during the economic crisis faced by the country. Since then, the TAC has been assigned in a precautionary way, by monthly increments, without information about the stock size and its distribution. Therefore, a more efficient and safe survey method that uses the currently available technology needs to be developed to achieve long-term monitoring of the resource. In this context, the present study shows and tests the performance and precision of Toki, a low-cost technology based on remote video imagery for scallop bed monitoring from small vessels. Specifically, this work evaluates the density and size structure of three scallop beds in San Jose Gulf. We study the accuracy of the Toki method for estimating the size structure compared to traditional diving and analyze the time required for all the previous processes.

## 2. Methods

### 2.1 Drop camera description and configuration

Toki is a pyramidal steel structure with a 100 cm × 100 cm square base and a height of 150 cm that holds two cameras on the top oriented to the base (**Figure 1**). The device’s dimensions were set to exactly include the1-m^2^ base of the quadropod within the video image. This structure (12 kg) is suitable to be maneuvered from a small semi-rigid boat. Toki has one high-definition (HD, **Figure 1** B-1) camera linked to the surface by a cable as part of a closed-circuit video (CCV) which is used for field operations. A second camera “slave” (Paralenz dive camera configurable up to 4K) records a copy of higher image quality which is used for data analysis (**Figure 1 B-2**). The CCV camera is connected to a 14-inch color monitor installed onboard through a class-5 FTP (foiled twisted pair), with an overall screen and supporting steel wire. The CCV camera is housed in a home-built waterproof compartment made of Grilon (polyamide alloys) with a 10-mm-thick acrylic lid, resistant to up to 15 atm. The CVC camera is used during fieldwork to correctly position the equipment on the bottom and avoid any type of collision or inconvenience, being able to monitor the entire process live from the surface. The video footage of the Paralenz camera (designed for underwater use, **Figure 1 B-2**) has a much better image quality and also records continuously a series of environmental variables (temperature and depth). Toki was deployed from a small semi-rigid boat (5.6 m). Two persons were required in addition to the boat driver: one to deploy the equipment manually and control the distance of the camera from the seabed and the other to real-time check Toki’s performance on the onboard monitor and to control video recording and register all necessary data (waypoint numbers, image quality, video timer, start and end time and waypoint points of each transect, etc.).

**Figure 1.**
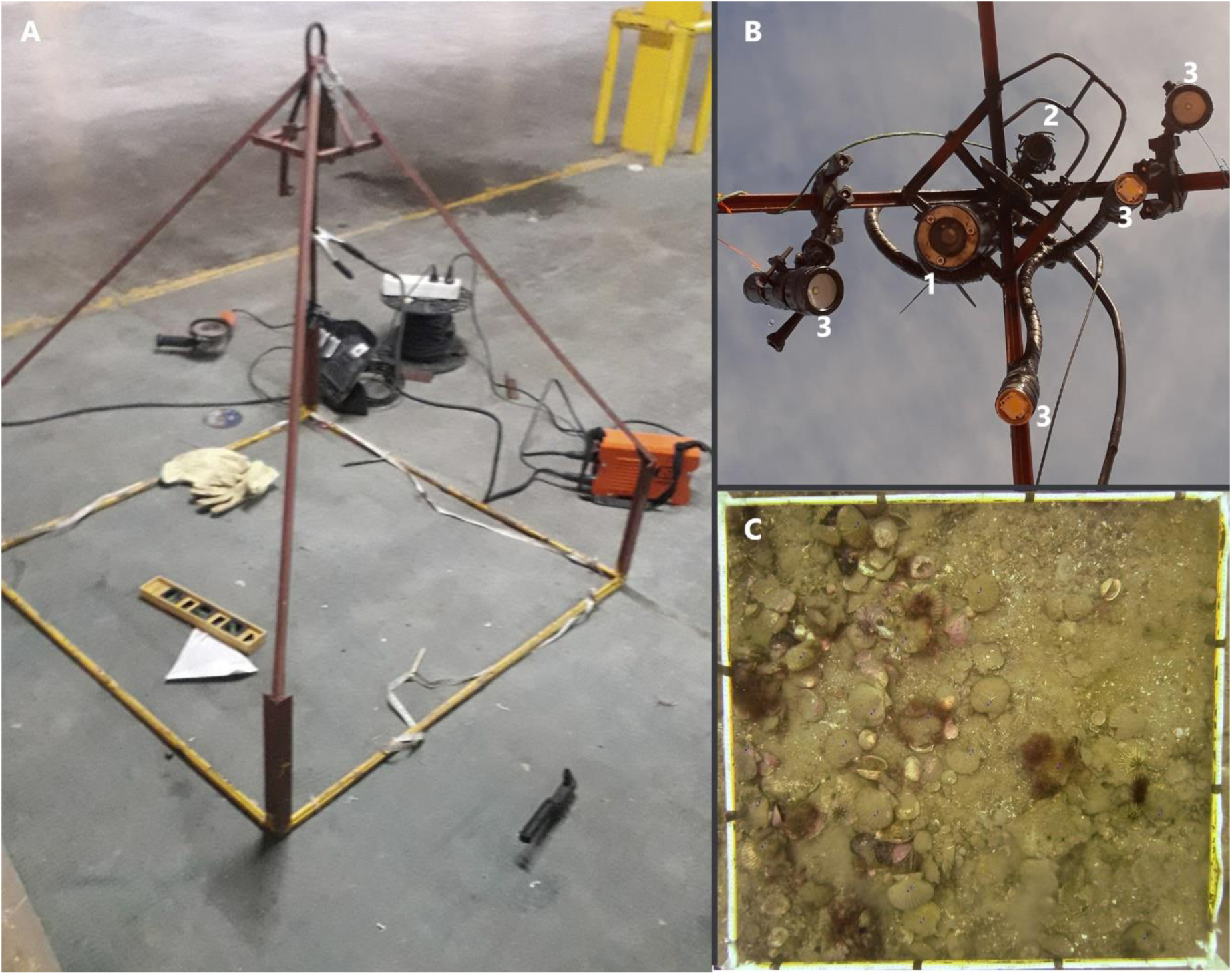
“A” Side view of the structure of the Toki video system, “B” Face-up view of the disposition of video cameras (B1= closed circuit camera; B2 = slave camera Paralenz; B3= video lights) and “C” snapshot (video-quadrat) of the slave camera view of a Toki.

**Figure 2.**
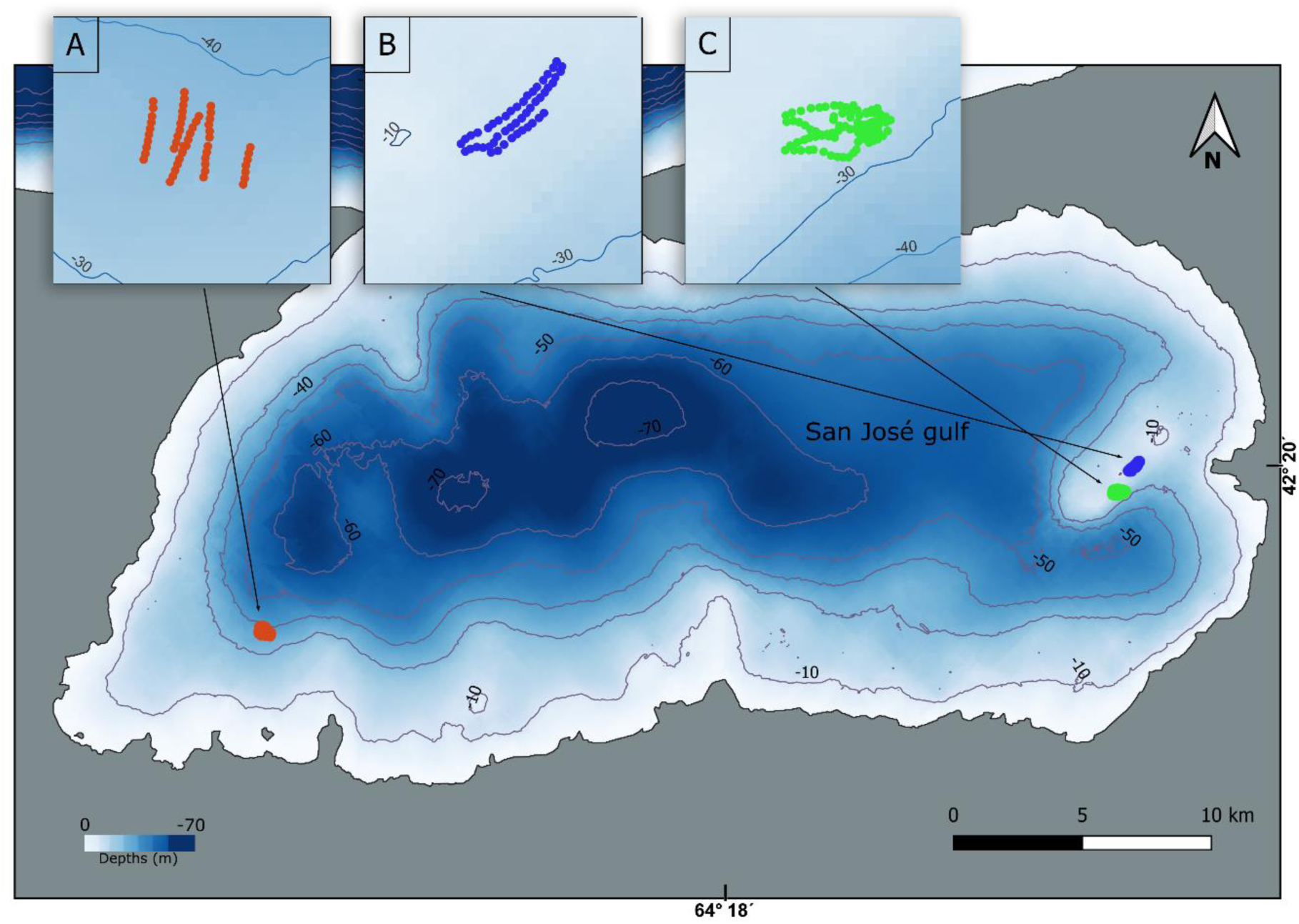
Location of the study area and the sampling sites at San José Gulf, North Patagonia: the three explored beds are indicated by the letters A, B and C, respectively.

### 2.2 Study site, experimental design and data collection

To evaluate Toki’s performance for estimating density and size composition of scallops in a natural bed, a study was conducted in SJG, Argentina (42.3°S, 64.3°W) in the spring and summer of 2021 and 2022, respectively (**Figure 2**). Three scallop beds were selected for this study. Bed “A” was located at about 40 m depth (below diving depth range) on the western sector of SJG which was discovered by the work team. The other two beds (“B” and “C”) were located off the eastern coast, between 12 to 40 m depth, on the slopes and foot of small seamounts (**Figure 2**). These last two beds were under exploitation during the 2021 season. Parallel video-transects were made on the three beds by towing Toki at low speed (3 knots maximum) and settling on the seabed every 30 m for approximately 8 seconds to take a snapshot (video-quadrat) from the video (“slave” camera) during each stop. The position (GPS waypoint) was recorded at each stop. Transects were conducted trying to cover a representative portion of the scallop bed and hence their length were not determined *a priori*.

### 2.3 Data processing

The video footage obtained by Toki was processed on a computer. The time of each settlement event is used for the snapshot extraction and to estimate its geographic position by synchronizing it with the GPS data (as described in Trobbiani *et al*., 2018) using an *ad-hoc* R script made for this purpose.

### 2.4 Estimation of density and size composition

The snapshots that included clear images of the seabed were used to count and estimate the size of scallops observed within the base of the quadropod (including those in contact with the borders). Counts provided a direct estimate of density (scallops per 1 m^2^). Three operators counted scallops on the entire set of images to study the variability between observers (1= AI, 2= GT and 3= LG). The first two were experienced in the use of video imagery for counts and the third had no experience or previous training.

To estimate the size-frequency distribution (SFD) all scallops that laid flat on the bottom and were perpendicular to the optical axis of the camera were measured from the umbo to the farthest ventral shell edge (shell height) using the free software ImageJ (https://imagej.nih.gov/ij/). Scallops tilted were not measured to avoid errors related to the lack of perpendicularity (Stobart *et al*., 2007; Willis and Babcock 2000). Only one operator (GT) took the measurements because the errors made in the process had been found to be negligible in a previous study (Trobbiani *et al*.,2018). A fixed reference scale included in the image (30 cm on the square frame; **Figure 1C**) allowed the software to scale size measurements. Blind measurements were directly stored in a text file for later construction of the SFDs for each bed.

### 2.5 Toki precision and accuracy assessment

A complementary study was conducted to evaluate the precision of scallop counts and SFDs obtained from video images versus the dive-based information. To that end a diver followed Toki underwater and after settlements of Toki, the diver collected all scallops found within the square frame (and in contact with the borders) and placed them in a separate mesh bag. A plastic number was shown to the camera before starting to collect the scallops and then placed into the mesh bag to allow assigning the sample to the correct video image. This protocol was repeated on 39 Toki settlements made on the shallowest zone of bed B (**Figure 2**). Collected scallops were counted and measured manually with a caliper, from the umbo to the furthest ventral shell edge, and data were compared to those obtained from the images, detailed previously (section 2.4).

The relative error of scallop counts from video images was modeled as a function of the following variables: depth (as a continuous variable), the true number of scallops collected by the diver in each square, the percentage of algal coverage (ranging from 0 to 100), and substrate type (Mud, Sand, Mud/Sand, Gravel, and Biogenic type). Mixed Effects Linear Models were employed using the *lme4* package 1.1-34 (Bates *et al.,* 2015) in the R software (R Development Core Team 2022). Initially, operator identity (three different operators counted the scallops) and quadrant identity were assessed as random effects. The Akaike Information Criterion (AIC) was used to compare the different adjusted models (Akaike, 1973), also implemented in R.

The percentage of algae coverage and substrate type variables were constructed using CoralNet (https://coralnet.ucsd.edu), an open-source resource for benthic image analysis. All snapshots taken by Toki were analyzed using 100 points within each image. The CATAMI classification scheme was employed to classify substrate types, aiming to adhere to a standardized vocabulary developed for identifying biota and benthic substrates from underwater images (see Althaus *et al*., 2015 for detailed information).

Differences between SFDs from Toki snapshots and by measuring scallops manually were evaluated with the Kolmogorov-Smirnov distance test. Using bootstrapping (n= 50,000), the Kolmogorov-Smirnov distances that would be obtained if both samples (video and shellfish) came from the same distribution were simulated.

### 2.6 Video imagery processing time

The time required to analyze the video imagery was recorded by the operators, including: the time spent counting scallops on each video-quadrat and the time of the processes of scaling snapshots and measuring scallops. The extraction of snapshots from the videos was done automatically, so it did not add any time. The total time used for estimating densities along the transects conducted in each of the banks was evaluated. In the case of the complementary study (section 2.5) the time spent estimating of sizes from snapshots by each of the operators was evaluated.

## 3. Results

A total of 17 video transects between 150 and 607 m in length (total length = 6491 m) were performed in the three beds at depths between 12 and 42 m in depth, requiring 375 min of total video time (**Table I**). In total, a polygon of ∼24 ha was covered (considered as the smallest polygon formed between the external points along transects).

**Table I.** Number of transects (**N transects**) made in each bed, total transects’ length (**Total length**) in meters, total time required to complete the transects in the field (**Time spent)** in minutes including travel time and other setbacks; (**Effective video time**) video time recorded in transects; total number of snapshots obtained for each bed (**N photo-quadrats**); for **Operator 1**, **2** and **3** shows the total (**N scallop counted**) for each one; (**Average density**) represents the scallop density averaged over the number of photo quadrats analyzed by each operator; (**SE**) standard error for each operator. Besides, the **Average density** and **mean scallop height** corresponding to sizes estimated by an operator 2 per bed are shown.

| Bed | N transects | Total length (m) | Time spent (min) | Effective video time | N photo-quadrats | Op. 1 |  |  | Op. 2 |  |  | Op. 3 |  |  |  | Mean scallop height |
| --- | --- | --- | --- | --- | --- | --- | --- | --- | --- | --- | --- | --- | --- | --- | --- | --- |
|  |  |  |  |  |  | N scallops counted | Average density | SE | N Scallops counted | Average density | SE | N Scallops counted | Average density | SE | Average density |  |
| A | 6 | 1592 | 165 | 104 | 53 | 195 | 3.68 | 0.71 | 252 | 4.75 | 0.69 | 258 | 4.87 | 0.79 | 4.4 | 71.7 |
| B | 4 | 1599 | 93 | 56 | 49 | 999 | 20.39 | 2.84 | 716 | 14.61 | 2.21 | 714 | 14.57 | 1.98 | 16.5 | 69.2 |
| C | 7 | 3300 | 117 | 110 | 101 | 1094 | 10.83 | 1.05 | 857 | 8.49 | 0.95 | 1065 | 10.54 | 1.14 | 10 | 68.1 |
| Total | 17 | 6491 | 375 | 270 | 203 | 2288 | 11.27 | 0.97 | 1825 | 8.99 | 0.77 | 2037 | 10.03 | 0.81 | 10.3 | 69.7 |

### 3.1. Density estimation

A total of 203 photo-quadrats (i.e., 203 m^2^ sampled) were extracted from the video imagery, all with adequate image quality to count and measure scallops (**Table I**).

On average, the three operators counted 2050 scallops, with densities ranging between 0 and 70 scallops/m^2^ (**Table I**). The mean relative errors ranged from 0.19 for operators 1 and 3 to -0.018 for operator 2.

### 3.2 Estimation of size frequency distribution

A total of 826 scallops laid perpendicular to the camera (i.e., Toki base) and were measured, representing about 45% of the total number of scallops counted by operator 2. Size frequency distributions differed among the beds, being unimodal in bed “A” and bimodal in the other two beds. The SFD in bed “A” had a mode at 75 mm (size media=71.7 mm; range= 52 to 90 mm). Bed “B” and “C” had similar SFDs ranging from about 45 to 94 mm, with a first mode at 57.5 and 59.7 mm, respectively, and a second mode at 70.6 mm and 76.1 mm. The largest scallop (94 mm) was registered in bed “C” (**Figure 3)**.

**Figure 3.**
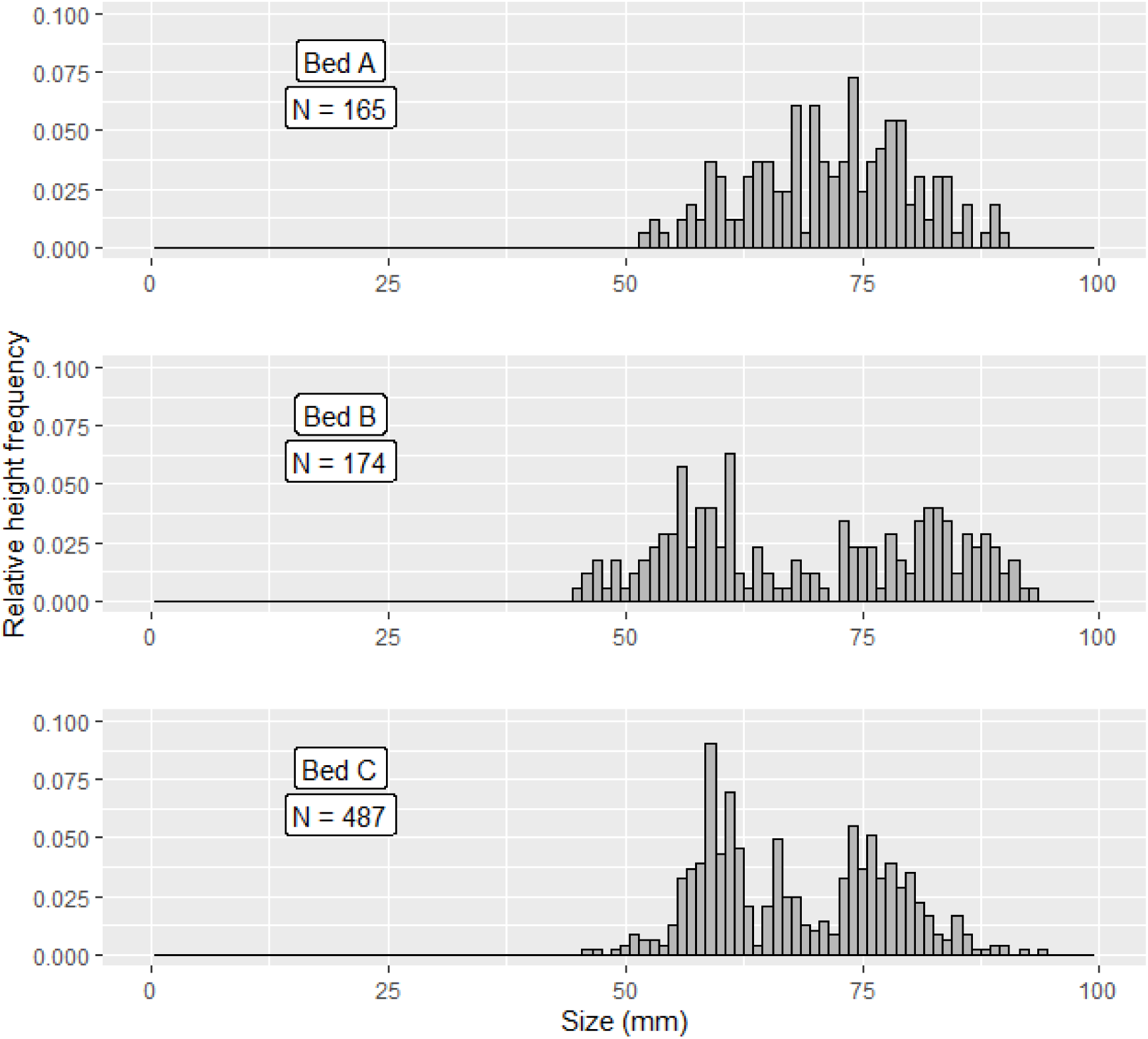
Relative height frequency estimated from the video snapshots for the three beds studied.

### 3.3. Accuracy and precision of scallop counts and size composition estimates

Divers collected 433 scallops (range 0 to 51 per quadrat) from 39 quadrats (1-m^2^) sampled in bed “B”. Corresponding scallop counts made by the three operators in the snapshots were 375, 366 and 262, indicating that they underestimated the total number of scallops encountered on the sampled video-quadrats by -13.4%, -15.5% and -39.5%, respectively. The mean relative errors were -33.3%, -21.3% and -37.8% for operators 1, 2 and 3, respectively. Densities estimated by operators 1 and 2 were more dispersed (sd=0.41 and sd=0.40) than those estimated by operator 2 (sd=0.29) (**Figure 4A)**.

**Figure 4.**
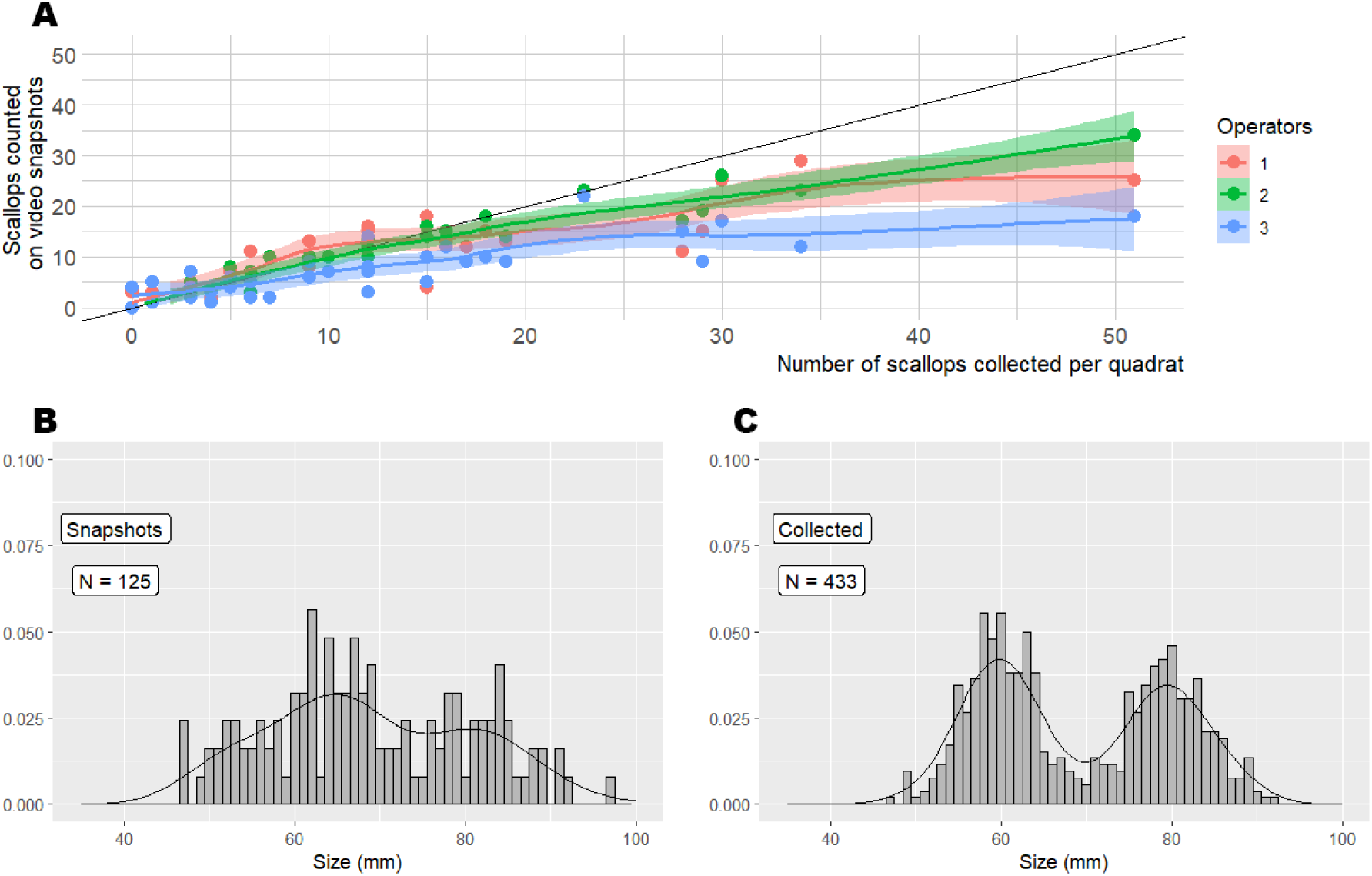
Estimation errors for scallop counts and size frequency distributions. **A:** video scallop counts made by three operators against the number of scallops collected by divers from each of the video snapshots. A smoothed curve fitted to the data from each operator is shown in different color. B and C: Relative size frequency distributions obtained from the video measurements (**B**) and from the sizes of the scallops collected from the same quadrats (**C**) in bed B.

The comparison of the full mixed-effect model (including all explanatory variables) with those that did not include random effects indicated that the variances contributed independently by the “Operator” and “ID-Photo”, and together, were not significant. The delta-AIC was 20.5 when contrasting the model without random effects against the best-fitting model with random effects (model with "Operator" and "ID-Photo"). Consequently, subsequent models were fitted using *lm*. Two models were preferred based on AIC and BIC: Model A, including the “true number of scallops” as the sole predictor, and Model B, which also included depth as a covariable. Model A showed a negative effect of the true number of scallops on the relative error, with an estimated coefficient equal to -0.0166 (p < 0.0001), and an adjusted R² of 0.1799. In Model B, incorporating both "depth" and "true number of scallops" the coefficient for depth was -0.0325 (p = 0.1102) and the coefficient for “true number of scallops” was -0.0149 (p < 0.0001), with an adjusted R² of 0.1928. Despite the lack of statistical significance for "depth"(p=0.1102), the increased R² indicates that is may have some contribution decreasing the error. The Akaike Information Criterion (AIC) favours Model B, as shown in table 2.

**Table 2.** Summary of the two selected *lm* models fitted to the errors of estimates count obtained, see the text for details. The structure for each model, parameter values, and the difference in AIC between the selected models are shown.

| ID | Model | Parameters values<br>(95% CI) | R<sup>2</sup><br>adjusted | Model<br>selection |  |
| --- | --- | --- | --- | --- | --- |
|  |  |  |  | df | ΔAIC |
| Model a | Relative error = intercept + real + $\varepsilon_i$;<br>$\varepsilon_i \sim N(0, \sigma_i^2)$; | intercept= 0.134 (0.058; 2.309)<br>“real”=-0.017 (0.003; -4.812) | 0.18 | 3 | 0.6 |
| Model b | Relative error = intercept + depth +<br>$\varepsilon_i$ + real + $\varepsilon_i$ ;<br>$\varepsilon_i \sim N(0, \sigma_i^2)$; | intercept= 0.727 (0.372;1.953)<br>“depth” = -0.033 (0.020; -1.612)<br>“real” = -0.015 (/0.004; -4.146) | 0.19 | 4 | 0 |

Of the 433 scallops collected by divers and measured to construct the SFD (**Figure 4c**), only 125 were correctly placed in the video quadrat image so that an unbiased shell height could be measured to construct the SFDs based on video quadrats (**Figure 4b**). The SFDs obtained with both methods presented a similar bimodal pattern, with one group of individuals at about 60 mm of shell height and second group at about 80 mm (**Figure 4b-c**). The Kolmogorov-Smirnov test, assessing the difference between the Survival Function Distributions (SFDs) of the ’video’ and ’ shellfish ’ methods, yielded a distance of d = 0.092. This calculated Kolmogorov-Smirnov distance did not reach statistical significance (p = 0.3611). The bootstrap analysis, comprising 50,000 replicates, further supported this conclusion, with the observed distance falling within a non-significant range of simulated distances.

### 3.5. Video imagery processing time

In total, the processing of video images (counting, scaling and measurement) took 15.5 hours. The processing time for the identification and count of scallops required an average of 110 min for the three beds, 102 min for 42 images from bed “A” (60, 53, 194 min for operators 1, 2, and 3, respectively), 75 min for 34 images from bed “B” (71, 65, 90 min for operators 1, 2 and 3, respectively), and 155 min for 73 images from bed “C” (133, 116, 216 min for operators 1, 2 and 3, respectively). Additional time used to set up the computer and pauses were not considered. The size measurements, taken only by operator 2, involved a total of 53 snapshots processed for bed “A”, where 165 scallops were measured in 71 minutes, 49 snapshots for bed “B” with 174 scallops measured in 126 minutes, and 101 snapshots for bed “C” with 487 scallops measured in 736 min. The time spent measuring scallops, which includes the scaling of each snapshot, took 53% more time on average than the identification and counting of live scallops. The total time required to scaling the snapshots, and counting and measuring each scallop was 1.12 min per scallop on average.

## 4. Discussion

Our results show that Toki is an efficient and accurate device, useful to estimate the density and size of sedentary or less mobile aquatic organisms lying close to the seabed. Particularly, results of present study shows an improvement in the quality of density data that can be obtained from Toki as compared with the underestimation biases introduced with the visual census during the traditional prospection protocol (Supplementary Material, Figure S1). Counts (i.e., density estimates) showed low levels of errors on experienced observers. However, higher errors occur in the case of the observer without experience on counts using video imagery. Our findings coincide with significant differences on levels of variability (not necessarily high levels of errors) found between experienced and non-experienced observers in previous studies involving visual estimation (e.g., Irigoyen *et al*., 2013, Kulbicky *et al*., 2010, Bernard *et al*., 2013). Operator training must be incorporated in the application of this method for monitoring.

The effects of covariate ("true number of scallops") analyzed on the level of the relative error on density estimates were significant. We observed a tendency to underestimate abundance in high-density areas (scallops ≥ 30 m2) and when the algal cover was greater than 15% (Trobbiani, personal observations). These two factors were associated in the study site (i.e., areas of high density presented red algae cover); hence, their effects could not be separated. Given that algae abundance presents seasonal variations, a simple way to mitigate the plausible effects of algal cover on the density estimations is to conduct surveys when algae are scarce. For example, *Undaria pinnatifida*, an alien alga in northern Patagonian gulfs, presents minimum abundance between March and June (Irigoyen., 2010; Casas et al., 2008; Dellatorre et al., 2012). In some beds with poor conditions, sampling by divers may be needed to avoid underestimation. Imperfect detectability (i.e., the inability to detect all individuals in a survey area) is one potentially important source of error that should be evaluated in all methodologies involving counts (Katsanevakis et al., 2012).

Size frequency distribution (SFD) obtained from the Toki video quadrats reproduced the actual SFDs obtained for the scallops collected by divers from the same quadrats evaluated by video imagery despite the small sample collected (N = 39) and the low fraction of scallops measurable per video-quadrat (∼ 25 % of the total present, collected by divers). The main advantage of using video imagery to estimate SFDs is that it can be used for beds located beyond safe diving depth range (e.g., > 25 m). Additionally, samples (i.e., video-quadrats) can be distributed over a larger spatial scale than what divers can achieve and hence may retrieve a more representative SFD of scallops beds for a comparable sampling effort (Trobbiani personal observations). The logistical costs associated with the use of video versus the work of experienced divers are minor, as well as the associated risks. Finally, the use of video imagery allows permanent recording for future analysis or data checks. However, when beds are in shallow waters, diving seems to be the most effective method to sample for size composition, given that taking measures from images is time-consuming (see Section 3.7). A trade-off between methods should be evaluated given specific objectives and logistical constrains.

Previous attempts to study Tehuelche scallops beds remotely using drift cameras (e.g., Fiorda *et al*., 2012) presented difficulties to estimate density given lack of spatial references to quantify area covered (Fiorda, personal observations). Video-quadrat methods similar to Toki have been also used to assess the stock of *Placopecten magellanicus* (Bethoney and Stokesbury, 2018, Stokesbury, 2002, Stokesbury *et al*., 2004, 2016). However, while similar in essence, the equipment used for sea scallops first heavy (450 kg) and was designed to be operated from oceanographic vessels with logistics and costs beyond the possibilities of most fisheries agencies in developing countries, especially when applied to small-scale fisheries. In this sense, the 12 kg of Toki and the fact that it can be operated from small boats are strengths of the method, making its implementation and transference possible. Additionally, the possibility of preserving the raw image data is relevant to future re-analyses of data and without the possible biases introduced by the observer (e.g., given by the change of the way in which distinct divers perform the same visual census).

Regarding the video imagery processing time reported in this study, time savings may be realistic through new developments using artificial intelligence. For example, the Coral Net software used in part of this work allows automatic annotations with just learning a few images. Using Toki in conjunction with these types of technologies could substantially improve its application.

Given the highly aggregated nature of scallops beds, the survey design used for Tehuelche scallops consisted of two stages. Over the last two decades, scallops beds monitoring on the study site has been conducted by visual census performed by divers within diving-safe depths (< 25 m) and complemented with in situ data collection (Parma *et al*., 2008; Soria *et al*., 2017). During the first stage, a diver towed from a boat counted and recorded all scallops observed over 1-m wide parallel transects laid perpendicularly to the coastline (5-25 m deep). While these counts have very large errors in dense patches, they are useful to locate and delimitate the scallop beds. In a second stage, density samples composed of ten 1-m2 quadrats distributed along the densest segment of each transect (were collected to correct the inherent bias of visual counts in high-density areas (supplementary material, Figure S1). The number of quadrats used to estimate density was limited by logistic and diving-time considerations. Toki could be used to replace the use of divers during both survey stages. In the first stage, Toki could be used as a drift camera to locate and delimitate the scallop beds defined by some threshold density (e.g., > 2 scallops/ m2). So, the first stage might be performed in a safer way including depths beyond the diving range and covering areas that are inaccessible with the current methodology. For the second stage, a larger number of video quadrats, covering the extension of the bed intercepted by each transect (normally from 100 to 1000 m), could be feasible given the easy operation of Toki. In summary, Toki could improve the data collection used to delimit scallop beds and estimate their average density, increasing the sample size and spatial coverage.

Altogether, Toki was used here to study the population structure and spatial distribution of the Tehuelche scallop stock but could be used similarly for other epibenthic species and also to map benthic habitats.

## 5. Conclusion

Assessment of Tehuelche scallops may be improved by the use of underwater video imagery. The Toki system provides a low-cost method for collecting information on density and size composition of scallop beds with minimal logistical field requirements. Further, survey improvements achievable with this technology may contribute to the sustainability of the artisanal diving fishery of Peninsula Valdés.

## Credit author statement

**Gaston Trobbiani**: Conceptualization, Methodology, Data acquisition, Data analysis Writing; **Alejo Irigoyen**: Conceptualization, Methodology, Data acquisition, Data analysis Writing, Supervision; **Leandro Getíno Mamet**: Data acquisition, Data analysis, Writing; **Ana Parma**: Conceptualization, Data analysis, Reviewing, Supervision.

## Acknowledgements

