## supplementary material, Figure S1 for "“Toki”, a light low-cost video system for seabed research: performance and precision of Tehuelche scallop (*Aequipecten tehuelchus*) survey estimates in San José Gulf, Argentina"

**Supplementary Material: Performance of Toki density estimation as compared with visual census during Tehuelche scallop prospection.**

To evaluate the Toki performance in the context of the Tehuelche scallop traditional prospection, the Toki estimation of density as a function of real number of scallops (this study data, Figure 4a) was compared with data from Tehuelche scallop prospection.

The Tehuelche scallop prospection work is divided into two stages. During the first the scallop beds are delineated, for which an artisanal shellfish-fisher diver is towed from a boat along transects and is responsible for counting the number of scallops observed over 100 m segments (visual census). In a second stage, and to avoid the bias inherent to visual counting in high density areas, the 100-m segments with scallop with counts >2 scallops/m$^2$ detected in the first stage are revisited. In each of these segments, ten 1m$^2$ squares are placed on the bottom and all scallops are collected to correct the density estimates from visual census. Then, a mean density is estimated for each segment from the quadrats data. As a result, visual census density and quadrat density data is available for each revisited high-density segment (Figure S1a). Given that between 1995 and 2006 the prospection protocol included 0.25m$^2$ (0.5m x 0.5m quadrat), the estimation were imprecise (Amoroso, 2012). For that reason, the Figure S1a was performed only with data from 2007 to 2017, when the used quadrat were of 1m$^2$, the same surface that Toki. More details of the scallop prospection methods in each year can be found in respective technical reports (Amoroso et al., 2010, 2011; Fiorda et al., 2013; Fiorda and Parma, 2015, 2012; Parma et al., 2008, 2007; Soria et al., 2017).

As well known for visual census, the prospection visual census data shows a great underestimation of density, which is progressive as scallop density increases (Figure S1a). While Toki is not exempt of this problem, the study shows a better performance than visual census performed by experienced shellfish-fisher divers during the scallop prospection (Figure 1b). This evidence an advance in the quality of the density data that can be obtained during the scallop prospection when using underwater cameras technology.

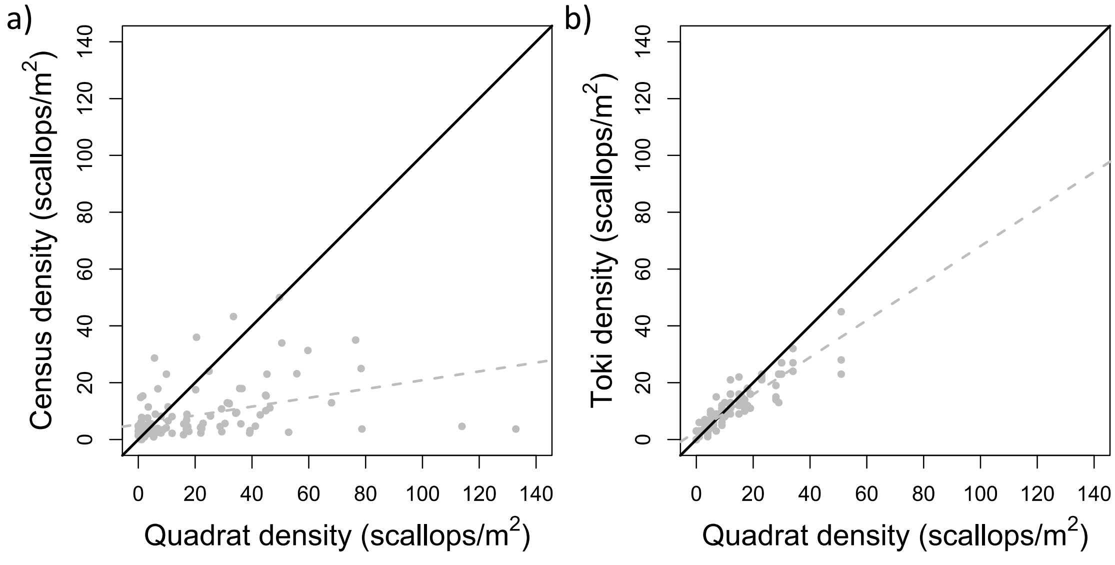


***Figure S1.*** *Estimations of scallop density. a) Data of visual census during the scallop prospection (2007-2017) in relation to in situ quadrat density data. b) Toki density estimation by three observers in relation to the quadrat scallop density estimated by diving collection of all scallops in the quadrat. Dashed line correspond to the adjusted regression rect.*
